# Macropinocytosis overcomes directional bias due to hydraulic resistance to enhance space exploration by dendritic cells

**DOI:** 10.1101/272682

**Authors:** Hélène D. Moreau, Carles Blanch-Mercader, Rafaele Attia, Zahraa Alraies, Mathieu Maurin, Philippe Bousso, Jean-François Joanny, Raphaël Voituriez, Matthieu Piel, Ana-Maria Lennon-Duménil

## Abstract

The migration of immune cells is guided by specific chemical signals, such as chemokine gradients. Their trajectories can also be diverted by physical cues and obstacles imposed by the cellular environment, such as topography, rigidity, adhesion, or hydraulic resistance. On the example of hydraulic resistance, it was shown that neutrophil preferentially follow paths of least resistance, a phenomenon referred to as barotaxis. We here combined quantitative imaging and physical modeling to show that barotaxis results from a force imbalance at the scale of the cell, which is amplified by the acto-myosin intrinsic polarization capacity. Strikingly, we found that macropinocytosis specifically confers to immature dendritic cells a unique capacity to overcome this physical bias by facilitating external fluid transport across the cell, thereby enhancing their space exploration capacity *in vivo* and promoting their tissue-patrolling function. Conversely, mature dendritic cells, which down-regulate macropinocytosis, were found to be sensitive to hydraulic resistance. Theoretical modeling suggested that barotaxis, which helps them avoid dead-ends, may accelerate their migration to lymph nodes, where they initiate adaptive immune responses. We conclude that the physical properties of the microenvironment of moving cells can introduce biases in their migratory behaviors but that specific active mechanisms such as macropinocytosis have emerged to diminish the influence of these biases, allowing motile cells to reach their final destination and efficiently fulfill their functions.

## Introduction

The adaptive immune system of mammals is composed of distinct cell populations that traffic between peripheral tissues and lymphoid organs. Adaptive immunity therefore relies on the ability of these cells to migrate all over the body, where they encounter numerous constraints imposed by the structure of the tissue (Heuze et al., 2013). Two important physical constraints that limit immune cell migration in tissues are geometrical confinement and hydraulic resistance (HR), which is induced by the interaction of a moving cell with the surrounding fluid (Bergert et al., 2015; Prentice-Mott et al., 2013). So far HR has been ignored as it exerts forces on cells that are supposed to be about a hundred fold smaller than the traction forces that cells themselves exert on the substratum (Bergert et al., 2015), suggesting that migrating cells should not be capable of sensing them. Nonetheless, recent findings highlighted that *in vitro* neutrophil-like cells preferentially migrate in confined environments towards low-hydraulic resistance paths (Prentice-Mott et al., 2013). This phenomenon, referred to as barotaxis, was attributed to the ability of these cells to sense and respond to differences in external hydraulic resistance using an active mechanism that may involve specific receptor(s) and signaling pathway(s). However, such mechanisms have not been identified so far. On the other hand, it was recently shown that cells migrating in confined environments use an adhesion-independent amoeboid-like migration mode that involves forces about a hundred fold smaller than the traction forces exerted by mesenchymal adhesive cells (Bergert et al., 2015). The magnitude of these forces is thus similar to those exerted by hydraulic resistance, suggesting that, in the absence of adhesions, hydraulic resistance could become the dominant resistive force that cells have to fight against to move.

We here combined theoretical physics, micro-fluidics and intravital imaging to investigate the mechanisms of barotaxis and its impact on immune cell behavior *in vivo*. We show that barotaxis results from a small force imbalance at the scale of the cell, which is amplified by the actomyosin network, while specific receptors and signaling pathways are not required. We further demonstrate that the ability of immature dendritic cells (DCs) to undergo macropinocytosis, i.e. non-specifically ingest extracellular fluid, renders them insensitive to HR and thereby facilitates their capacity to patrol their environment. By contrast, mature DCs are no longer macropinocytic and may benefit from barotaxis to find low-resistance paths while migrating to lymph nodes to initiate the immune response. Thus, specific mechanisms have emerged for DCs to overcome the physical obstacles they are exposed to *in vivo* and efficiently achieve their immune sentinel function.

## Results and discussion

### A theoretical framework to predict barotaxis

As mentioned above, the range of forces exerted by non-adherent cells migrating under confinement (such as immune cells) is comparable to the range of HR forces. Therefore, we hypothesized that barotaxis could result from a purely mechanical interaction of the cell migratory apparatus with the surrounding fluid. In other words, cells would preferentially migrate to lower HR paths because these paths oppose the lowest force to their migration. To test this hypothesis, we extended a well-established hydrodynamic physical model that successfully reproduces the fast mode of migration of non-adherent cells in confinement from a minimal set of hypotheses (Callan-Jones and Voituriez, 2013). In this model, cells are described as made of an active poro-elastic material, whose constitutive equations stem from the symmetries and conservation laws of the cortical actomyosin system (Joanny et al., 2007; Juelicher et al., 2007; Kruse et al., 2005). The main parameters of the model are: (1) the amplitude of the contractile stress, which encodes the interaction between myosin-molecular motors and the cortical actin and (2) the cell membrane-fluid permeability, which encodes the forces arising as the extracellular fluid passes through the confined cell. The model harbors a simple motility mechanism in 1d-confined environments that is critically controlled by both the level of contractility and the cell length. Above a critical level of activity, cells exhibit a spontaneous polarization mechanism: random fluctuations of 1d non-polarized cells are amplified, breaking the front-rear symmetry through an accumulation of contractile activity at the back. This drives spontaneous retrograde flows of the actin cortex, resulting in a net thrust force on the channel walls and therefore a polarized motile state. To investigate the mechanism of barotaxis, we extended this model (Callan-Jones and Voituriez, 2013) from a 1d-channel to a three-way bifurcation (See Supplementary Information) exhibiting different values of HR in the 2 upper paths (**Fig. 1A**). Barotaxis was quantified in these three-way bifurcations as the bias of cell direction toward lower resistance path. We observed that this directionality bias increased progressively with HR asymmetry until it reaches an upper threshold of 0.45 (corresponding to about 70% of cells choosing the low resistance side) (**Fig. 1B**), indicating that our schematic model is sufficient to recapitulate barotaxis. Therefore, barotaxis can be predicted by the mechanical interaction between the migrating cell and its environment without the need to introduce any receptor or signaling pathway dedicated to pressure sensing.

**Figure 1:**
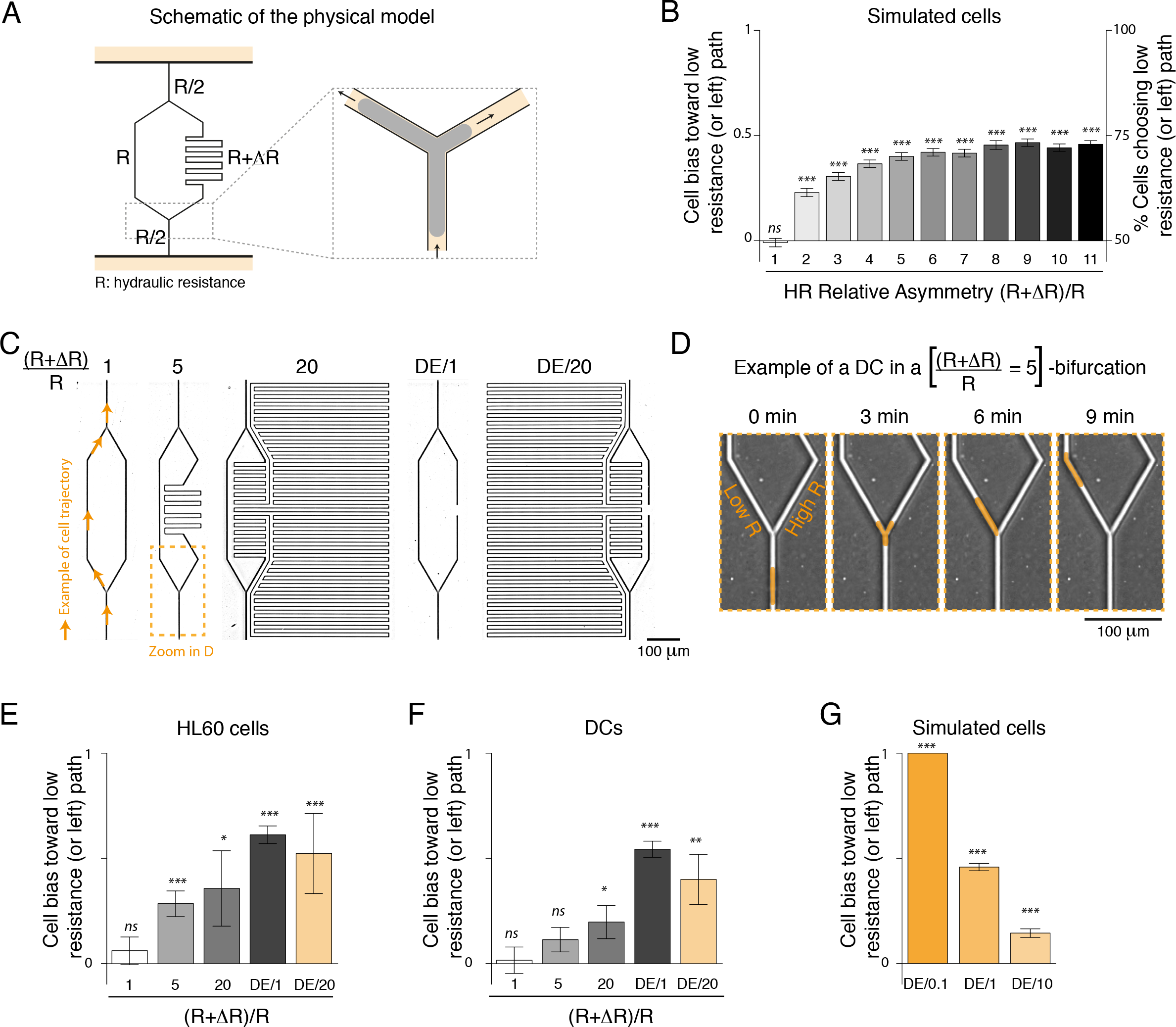
Barotaxis results from force imbalance. A. Schematic representation of the physical model. Grey: simulated cell. Light orange: surrounding fluid. B. Bias of simulated cells toward the low resistance path. Bias is computed as (N_LR_−N_HR_)/(N_LR_+N_HR_), where N_LR_ and N_HR_ are the number of cells choosing low and high resistance paths resp., meaning that 0 corresponds to no bias (50% of cells on each side), 1 corresponds to full bias (100% of cells choosing low resistance path). Percentage of cells choosing low resistance path is indicated on the right axis for reference. C. Transmitted light images (inverted LUT) of microfluidic chips exhibiting different hydraulic resistance on both arms of the bifurcation. Calculated values of HR relative asymmetry are indicated above each bifurcation type. D. Example of a dendritic cell migrating in a bifurcation ((R+ΔR)/R=5) and choosing low resistance path. Cell has been highlighted in light orange for clarity. E. Bias of HL60 cells in the different microbifurcations. E. Bias of DCs in the different microbifurcations. F. Bias of simulated cells in bifurcations exhibiting a dead-end versus a high resistance path. Data are pooled from at least 4 independent experiments. Statistics above graph bars indicate difference to the value 0 (corresponding to no bias) calculated by a one sample t-test. ***: p-value < 0.001; **: p-value < 0.01; *: p-value < 0.05; *ns*: non significant.

To experimentally address the predictions inferred from the model, we designed Y-shaped micro-fabricated devices exhibiting different HR and a small cross-section (18µm^2^) to ensure confinement (**Fig. 1C**). We verified that HR asymmetry did not induce any asymmetry in terms of channel wall coating or medium filling (**Fig. S1**). We let neutrophil cell lines (HL60) as well as immature dendritic cells (iDCs) migrate in the different bifurcations and quantified their bias toward the lower resistance side as we did in the cell simulation (**Fig. 1D** and **movie S1**). In accordance with the model prediction, we observed a gradual response to HR of both HL60 (**Fig. 1E**, consistent with published data (Prentice-Mott et al., 2013) and iDCs (**Fig. 1F**) up to a certain threshold of 0.61 and 0. 54 resp. (corresponding to about 80% of cells choosing the low resistance path). Neutrophil-like cells were slightly more sensitive to low increments in HR (×5 and ×20) as compared to iDCs. Noticeably, a bias of 1, which corresponds to 100% of cells choosing the low resistance path, was never observed (even when cells faced an infinite HR generated by a dead-end (bifurcation DE/1)). In order to test whether cells were responding to an absolute value of HR or a relative one, we modified our dead-end bifurcation (DE/1) to increase the HR of the low resistance side (bifurcation DE/20, **Fig. 1C**). In these bifurcations, we observed the slightly decreased bias (**Fig. 1E-F**) predicted by our model (**Fig. 1G**). These experimental and theoretical results suggest that cells are sensitive to differences in HR and choose the side with a relatively lower resistance, with a sensitivity that only weakly depends on the absolute HR value. Thus, the experimental data obtained with confined HL60 and iDCs agrees with the predictions obtained from the simulations and validate our hypothesis: barotaxis is a mechanical process in which cells choose the path opposing the least resistance to migration.

### A small force imbalance amplified by the actomyosin network helps cells choosing low-resistance paths

In order to better understand how cells respond to HR asymmetry and choose a direction, we characterized the shape dynamics and actin distribution of simulated cells (**Fig. 2A** and **movie S2**) and compared it to iDCs facing bifurcations (**Fig. 2B** and **movie S2**). We focused on symmetric (X1) and fully asymmetric (DE/1) bifurcations. The dynamics predicted by the model can be interpreted as follows: at the bifurcation, cells form two extending arms (one per outlet). The subsystem constituted by the two arms eventually reaches a critical size and its actomyosin network self-polarizes spontaneously towards one of the two available directions through a contractile instability analogous to the spontaneous polarization mechanism observed in 1d channels (Callan-Jones and Voituriez, 2013). This leads to the retraction of the arm with increased actin content and thus the choice of a direction (**Fig. 2C**). Strikingly, even in asymmetric bifurcations, both arms extend symmetrically, independently of HR values, and only a small difference in extension could be observed prior to the directional choice (**Fig. 2C**). In the model, this effect results intrinsically from the fact that the choice of direction is made through a dynamic instability of the actomyosin cytoskeleton. When the total spatial extension of the subsystem constituted by the two upper arms is smaller than a critical length, this subsystem is stable and only weakly responds to the hydraulic resistance bias. When reaching the critical length, this upper subsystem becomes unstable, its actomyosin network self-polarizes and dramatically amplifies any infinitesimal difference existing between the extensional speeds of the two arms. In symmetric bifurcations, this difference is exclusively due to fluctuations and the resulting cell direction is thus random. In contrast, in asymmetric bifurcations, the arm extensional speed is on average slightly faster toward the path of least resistance. The amplification of such small difference therefore leads to a significant barotactic bias toward this path (See Supplementary Information). These non-trivial predictions of the model were confirmed experimentally by quantifying arm extension in iDCs. When encountering a bifurcation, cells extended two arms (one in each microchannel path) until one arm retracted, resulting in a choice of direction (**Fig. 2B**). Strikingly, both arms extended at very similar speeds, even in asymmetric bifurcations (**Fig. 2D** and **S2**). This experimental observation strongly suggests that barotaxis is well described by the model: the difference in mechanical load stemming from the difference in the HR of the two paths slightly slows down the arm facing the highest resistance path; this narrow difference is then amplified by the spontaneous polarization of the actomyosin network, leading to a strong bias in the directional choice.

To obtain direct experimental evidence for the role played by the actomyosin system in this amplification mechanism, we quantified the distribution of actin in simulated cells and LifeAct-GFP transgenic iDCs. In both symmetric and asymmetric bifurcations, we observed that retraction of the losing arm was associated to actin accumulation (**Fig. 2E** and **S2**). This accumulation was observed earlier when cells were facing asymmetric bifurcation compared to symmetric bifurcations (Fig. 2E, **right**), consistent with the model’s predictions (Fig. 2E, **left**).

**Figure 2:**
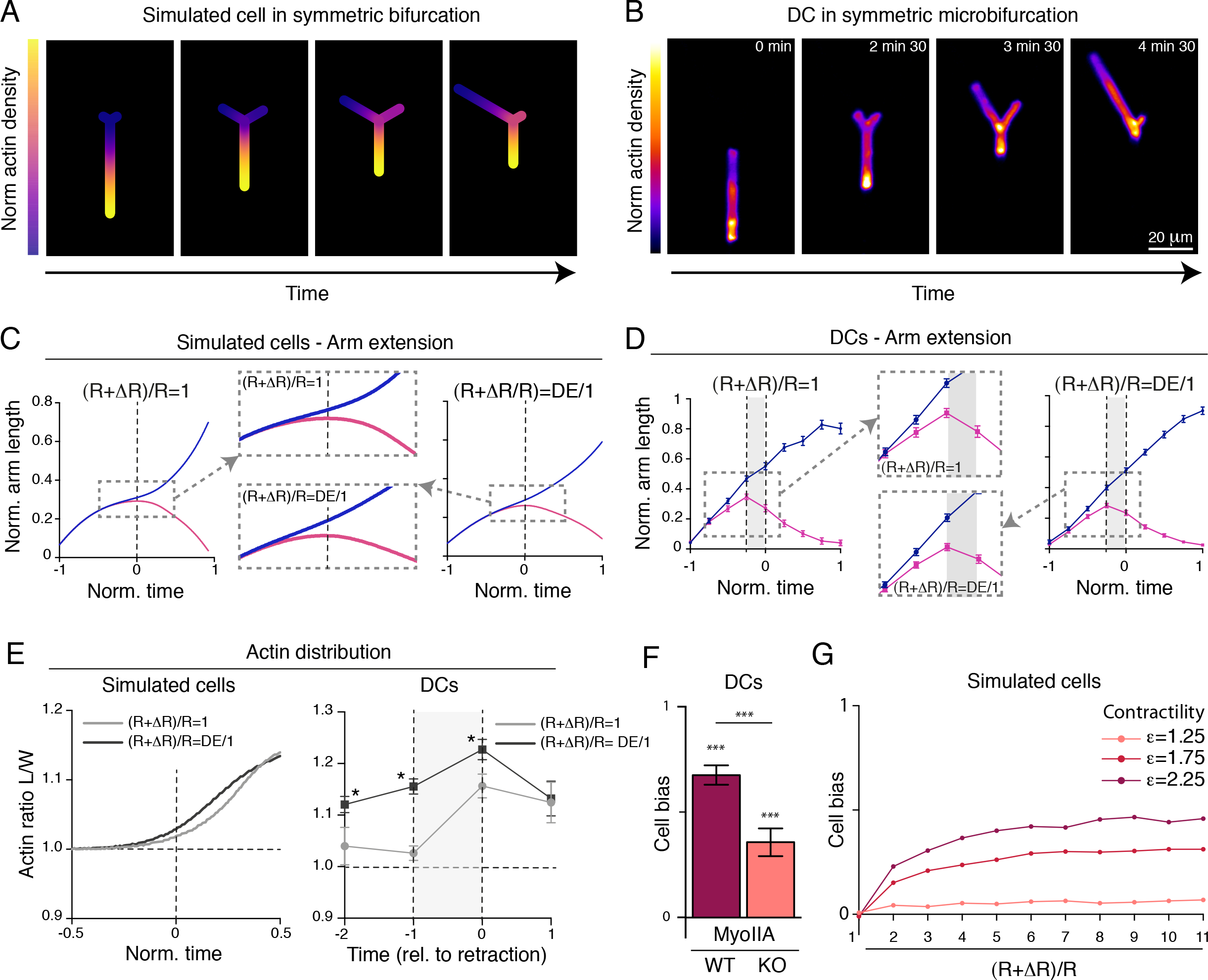
Directional choice is driven by spontaneous polarization of actomyosin cytoskeleton. A. Example of actin distribution of a simulated cell in a symmetric bifurcation. B. Example of LifeAct distribution of a DC in a symmetric bifurcation. C. Normalized arms length of simulated cells is plotted against normalized time (centered around retraction time). D. Average normalized arm length of DCs is plotted against normalized time (centered around retraction time). E. Average actin ratio between losing and winning arms for simulated cells and DCs are plotted against time. Statistics are calculated by a t-test. F-G. Barotaxis increases with contractility. F. MyoII-WT and MyoII-KO DC bias in DE/1 bifurcations. G. Simulated cell bias for different values of contractility normalized by the value at the onset of the instability. Statistics above graph bars indicate difference to the value 0 (corresponding to no bias) calculated by a one sample t-test. Statistics between graph bars indicate difference between conditions calculated by a Mann-Withney test. ***: p-value < 0.001; **: p-value < 0.01; *: p-value < 0.05; ns: non significant. Data are representative of at least 3 independent experiments.

These findings support the idea that the actomyosin network does indeed spontaneously polarize and directly controls arm retraction and thus the choice of direction. Accordingly, we experimentally found that impairment of actomyosin contractility (in Myosin II knock out (KO) iDCs) decreased the directional bias toward the low resistance path (**Fig. 2F**), as predicted by the model (**Fig. 2G**). Our findings thus demonstrate that HR asymmetry strongly biases cell directionality by generating a small force imbalance that is further amplified by the actomyosin cytoskeleton. Therefore, the intrinsic properties of the actomyosin system are sufficient to explain barotaxis, with no requirement for specific pressure sensors.

### Macropinocytosis provides immature DCs with a unique capacity to overcome barotaxis

We reasoned that an important parameter for barotaxis should be the permeability of the cell to the surrounding fluid, which is quantified by the cell-fluid resistance parameter ζ. If cells oppose a strong resistance to permeation flows (low permeability, large ζ), the forces experienced by the two cell arms at a bifurcation are controlled by the difference in hydraulic resistance, and could be in principle infinite in the case of a dead-end. On the other hand, if fluid flows freely through the cell (large permeability, low ζ), migration can occur without moving the extracellular fluid and hydraulic resistance is irrelevant. To test this parameter, we quantified barotaxis in our model for different levels of cell permeability (obtained by varying ζ/R). We consistently found that cells endowed with a large effective permeability (ζ/R=10^−3^) are insensitive to barotaxis (bias of 0), while cells exhibiting a small effective permeability (ζ/R=10^−1^) are extremely sensitive to HR asymmetry (**Fig. 3A**). Therefore, our experimental results showing that cells have a strong bias towards open ends, but that the bias never reaches 100% (**Fig. 1**), suggest that they are slightly permeable but still oppose a resistance to fluid flow.

**Figure 3:**
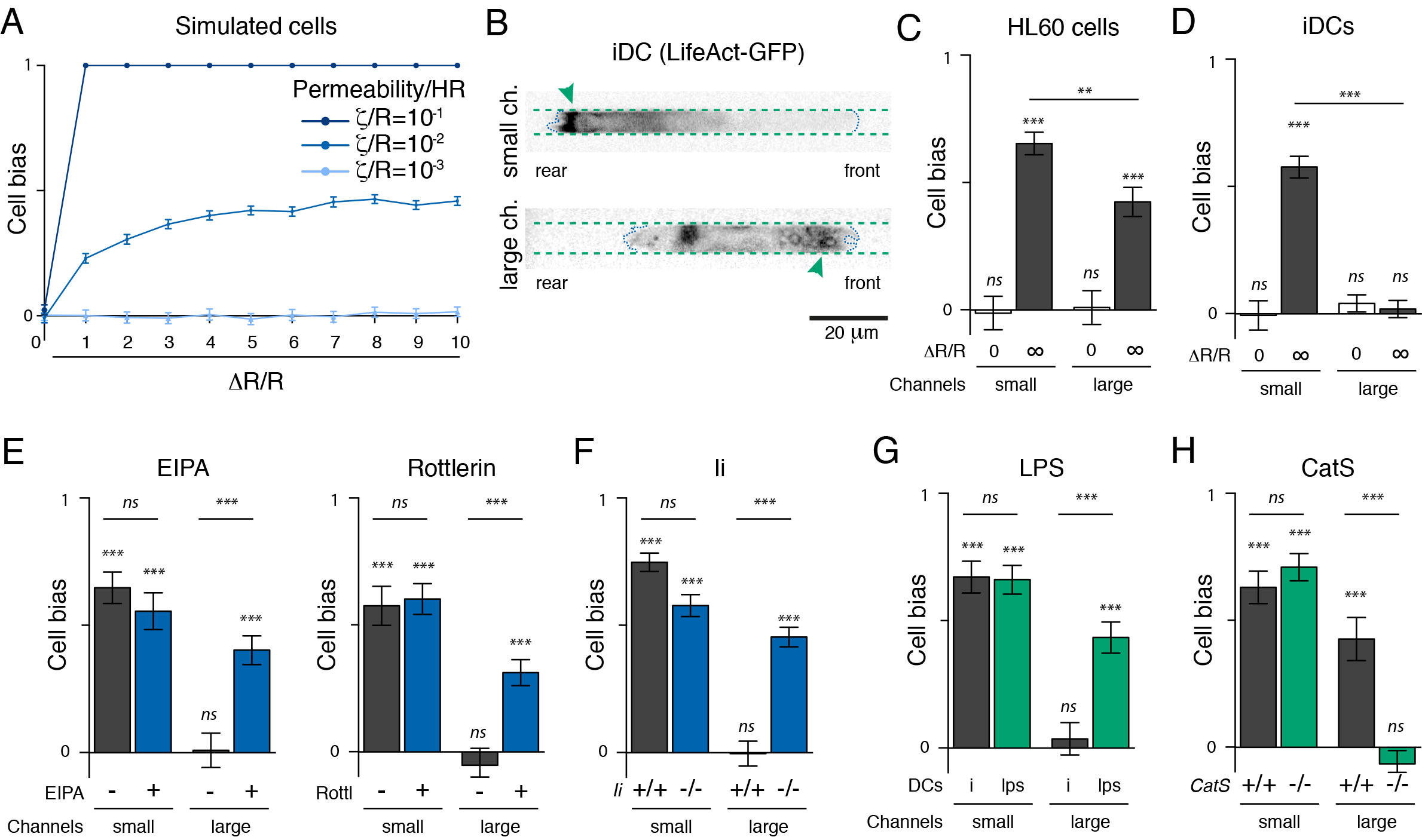
Macropinocytosis increases dendritic cell permeability to the extracellular fluid and therefore abolishes barotaxis. A. Barotaxis decreases with cell permeability. Bias of simulated cells for different values of the cell permeability to hydraulic resistance ratio. B. iDCs do macropinocytosis in large channels but not in smalls. Example of LifeAct images (inverted LUT). Channel walls are indicated by a green dotted line. Cell front and rear are highlighted by a blue dotted line. Small channel: green arrow indicates actin accumulation at the rear. Large channel: green arrow points to macropinosomes. C. Bias of HL60 (left) and iDCs (right) toward low resistance path in small and large DE/1 microchannels are graphed. D. Inhibiting macropinocytosis restores barotaxis in large channels. Bias of iDCs treated or not with EIPA (left) or Rottlerin (center), or deficient for Ii (right) in DE/1 channels are plotted. E. Mature DCs are biased in large channels. Bias of DCs treated or not with LPS are graphed. F. Bias of mDCs deficient or not for CatS in DE/1 channels are plotted. Statistics above graph bars indicate difference to the value 0 (corresponding to no bias) calculated by a one sample t-test. Statistics between graph bars indicate difference between conditions calculated by a Mann-Withney test. ***: p-value < 0.001; **: p-value < 0.01; *: p-value < 0.05; ns: non significant. Data are pooled from at least 3 independent experiments.

These observations, together with the predictions of the model, made us wonder whether some specific cells could be permeable enough to exhibit no bias at all and freely explore dead-ends, similarly to the largely permeable simulated cells (ζ/R=10^−3^, **Fig. 3A**). Cells can be permeable to fluid by leaving small interstices between the cell body and the channel walls, or by taking up fluid at their front and releasing it at their back. Different mechanisms can be at stake: (1) fluid can enter and move passively out of the cell using membrane channels such as Aquaporins or (2) fluid can be taken up and secreted via vesicles. The latter is particularly efficient in iDCs as they are endowed with the ability to constitutively internalize extracellular fluid through macropinocytosis (Sallusto et al., 1995). This evolutionary conserved mechanism relies on the formation of giant vesicles (> 200 nm) where liquid is internalized in a non-specific manner (Buckley and King, 2017). We thus hypothesized that iDCs performing macropinocytosis may not be barotactic. To test this hypothesis, we manipulated the macropinocytic capacity of iDCs using different strategies. First, we modified the dimensions of Y-shaped microchannels to allow for efficient formation of macropinosomes. Indeed, in small channels as the ones used in Fig. 1 and 2 (< 20 um^2^-section), F-actin is barely recruited to the cell front (our unpublished data) and therefore macropinosomes do not form, contrary to what had been observed in larger channels (> 30 um^2^-section (Chabaud et al., 2015)) (**Fig. 3B**). We thus quantified the bias of HL60 cells and iDCs in symmetric (X1) and asymmetric (DE/1) bifurcations of both small (18 µm^2^) and large (32 µm^2^) cross-sections. HL60 cells, which are known as non-macropinocytic cells, exhibited no bias in symmetric bifurcations, but were strongly barotactic when facing dead-ends (**Fig. 3C**). Yet, we observed a small decrease in barotaxis in large channels compared to small channels. This can probably be explained by the lower level of confinement of HL60 cells in the large channels, which could let fluid flow freely on the sides of the smaller cells. iDCs are bigger than HL60 cells, and are thus well confined both in small and large channels (**Fig. 3B**). In small channels, iDCs had most of their actin localized at the back, and did not perform any giant vesicle, suggesting they are not macropinocytic in these conditions (**Fig. 3B**, **top**). As expected, there were biased toward low resistance path when facing asymmetric bifurcations (**Fig. 3D**). However, in large channels, we could observe large vesicles covered in actin at the front of the DCs (Fig. 3B, **bottom**), suggesting they do perform macropinocytosis, and as predicted by the model, they totally lost barotaxis and explored dead-ends equally well as open-ends (**Fig. 3D**). Consistent with these data, pharmacological inhibition of macropinocytosis using EIPA (also known as Amiloride) or Rottlerin, two specific macropinocytosis inhibitors described in the literature (Koivusalo et al., 2010; Sarkar et al., 2005; West et al., 1989), restored barotaxis in iDCs migrating in large microchannels (**Fig. 3E**). These inhibitors had no impact on cells migrating in small channels (**Fig. 3E**). Equivalent results were obtained when genetically compromising macropinocytosis by knocking out (KO) the gene encoding for CD74/Ii, which recruits Myosin II at the front of DCs to promote macropinosome formation (Chabaud et al., 2015): Ii KO iDCs were non macropinocytic and barotactic even in large channels (**Fig. 3F**). We further confirmed these findings by indirect inhibition of macropinocytosis (Vargas et al., 2016) using the Arp2/3 inhibitor CK666 (**Fig. S3**). Altogether these results show that, as predicted by our model, permeability to external fluid is a key parameter for barotaxis. They demonstrate that macropinocytosis can efficiently suppress the directional bias introduced by HR, likely by increasing cell permeability to extracellular fluid.

### Down regulation of macropinocytosis in mature DCs restores barotaxis

When iDCs are activated by danger-associated signals (such as microbial components), they enter into a “maturation” program that triggers their migration to lymphatic vessels and lymph nodes where they activate T lymphocytes to launch the adaptive immune response. Importantly, macropinocytosis is down regulated during this maturation process (Sallusto et al., 1995). Accordingly, we found that treatment of iDCs with the bacterial wall component lipopolysaccharide (LPS) was sufficient to inhibit macropinocytosis and promote barotaxis even in large channels (**Fig. 3G**). DCs KO for the protease Cathepsin S (CatS), which remain macropinocytic upon LPS treatment (Chabaud et al., 2015), were not barotactic in large channels (**Fig. 3H**). This was also confirmed by treating mature DCs (mDCs) with the formin inhibitor Smifh2: as previously shown (Vargas et al., 2016), this small molecule impaired macropinocytosis down-regulation and accordingly, it compromised barotaxis (**Fig. S3**). Hence, the down-regulation of macropinocytosis that accompanies DC maturation is associated to the acquisition of barotactic properties.

### Macropinocytosis increases with hydraulic resistance

Our results showing that macropinocytosis allows iDCs overcoming HR and exploring dead-ends prompted us to closely monitor this process before and after cells chose their direction. For this, we followed in time the distribution of LifeAct-GFP, as we have shown that actin accumulation at the front of iDCs can be used to quantify macropinocytosis (Vargas et al., 2016). Interestingly, we found that in small channels, which diminish the macropinocytic process, the few cells that showed actin accumulation at their front before reaching the bifurcation (**Fig. S4**) chose open-end at 51% and dead-end at 49%, thus rendering no bias. In contrast, cells showing no actin accumulation at their front before reaching the bifurcation chose open-ends at 78%. Hence, even in small channels, the few cells that displayed characteristics of macropinocytosis (front actin) are insensitive to HR whereas non-macropinocytic cells, including mDCs, strongly undergo barotaxis.

When analyzing the few iDCs that chose dead-ends in these channels, we observed that most of them reached the dead-end, suggesting that they were able to cope with the amount of fluid that was contained in the channel. Remarkably, we noticed that they actively extended and retracted their front and ingested increased amounts of extracellular fluid compared to the iDCs choosing open-ends (**Fig. 4A-B**). Interestingly, this change of behavior was also observed when comparing iDCs migrating in dead-ends versus open-ends in large channels (**Fig. 4A-C**). We therefore hypothesized that elevated HR stimulates the macropinocytic process itself, thereby helping cells cope with the external fluid while migrating in high-resistance paths. Signatures of the macropinocytic activity include a decrease in cell speed and an increase in the protrusion/retraction activity in addition to actin accumulation at the cell front (Chabaud et al., 2015; Vargas et al., 2016). Strikingly, all these hallmarks of macropinocytic cells were observed in dead-ends but not in open-ends in small channels, and were enhanced in dead-end large channels compared to open-end counterparts (Fig 4D-F **and S4**). These observations indicate that iDCs react to changes in HR by increasing their capacity to ingest extracellular fluid, suggesting that macropinocytosis can be modulated by environmental physical cues.

**Figure 4:**
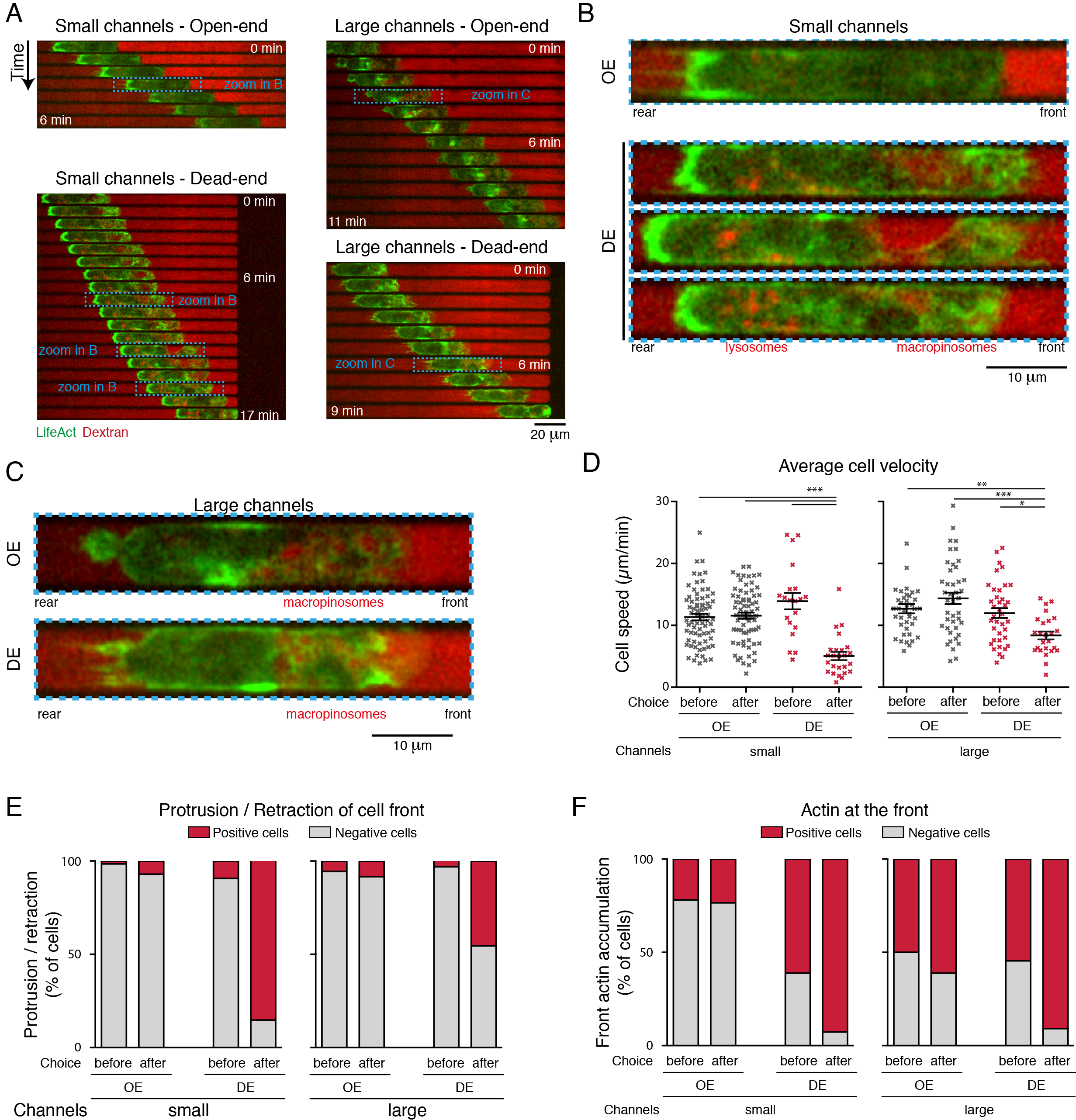
High hydraulic resistance enforces a macropinocytosis like behavior in dendritic cells. Cells have been classified depending on their directional choice toward open (OE)-or dead-end (DE). A-C. Time lapse images of iDCs in small or large channels, after they have chosen open-or dead-end sides. Green: LifeAct. Red: Dextran-AlexaFluor647. Macropinosomes and lysosomal regions are indicated when visible. D. Average cell speed. One dot corresponds to one cell. Statistical analysis was performed using a one-way ANOVA. E. Protrusion / retraction activity. Cells are classified as positive for protrusion / retraction activity if at least one protrusion / retraction of the front is clearly observed in the last 2 min before it reached bifurcation (“before”) or during the 3 minutes after it exited the bifurcations (“after”). F. Front actin. Cells are classified as positive for actin accumulation at their front if actin accumulation is clearly visible at least once in the last 2 min before it reached bifurcation (“before”) or during the 3 minutes after it exited the bifurcations (“after”).

### Macropinocytosis favors space exploration while barotaxis enhances directionality in complex environments

The lack of sensitivity of iDCs to HR might improve their capacity to explore tissues in an unbiased way, making them more efficient as immune sentinels. Therefore, macropinocytosis, which is used by iDCs to sample tissues for the presence of danger-associated molecules-for example those belonging to infectious agents-a key step in the initiation of immune responses, could also contribute to the patrolling function of iDCs by allowing them to overcome barotaxis and thus explore spaces inaccessible to other cells such as dead-ends.

In order to investigate the role of macropinocytosis in tissue exploration *in vivo*, we generated mixed bone marrow chimeric mice containing both macropinocytic (wild-type, WT) and non-macropinocytic (Ii KO, (Chabaud et al., 2015)) iDCs that can be distinguished based on the fluorescent protein (GFP or YFP, resp.) they express. The migration of iDCs in the ear skin was assessed by two-photon imaging, at steady state or after induction of an edema by sub-cutaneous injection of λ-carrageenan (Winter et al., 1962) (**Fig. 5A**). Space exploration was quantified as the volume explored by Ii WT or Ii KO cells over time (**Fig. 5B**). At steady state, similar volumes were explored by macropinocytic DCs (WT) and non-macropinocytic DCs (Ii KO). In contrast, macropinocytic DCs (WT) were significantly more exploratory than their non-macropinocytic counterparts (Ii KO) in the presence of an edema (**Fig. 5C**). Hence, cells permeable to fluid, and thus insensitive to HR, such as macropinocytic immature DCs, have an increased capacity to explore inflamed tissues in which the volume of fluid to displace is higher. This might help iDCs to uphold their sampling function in inflamed tissues.

**Figure 5:**
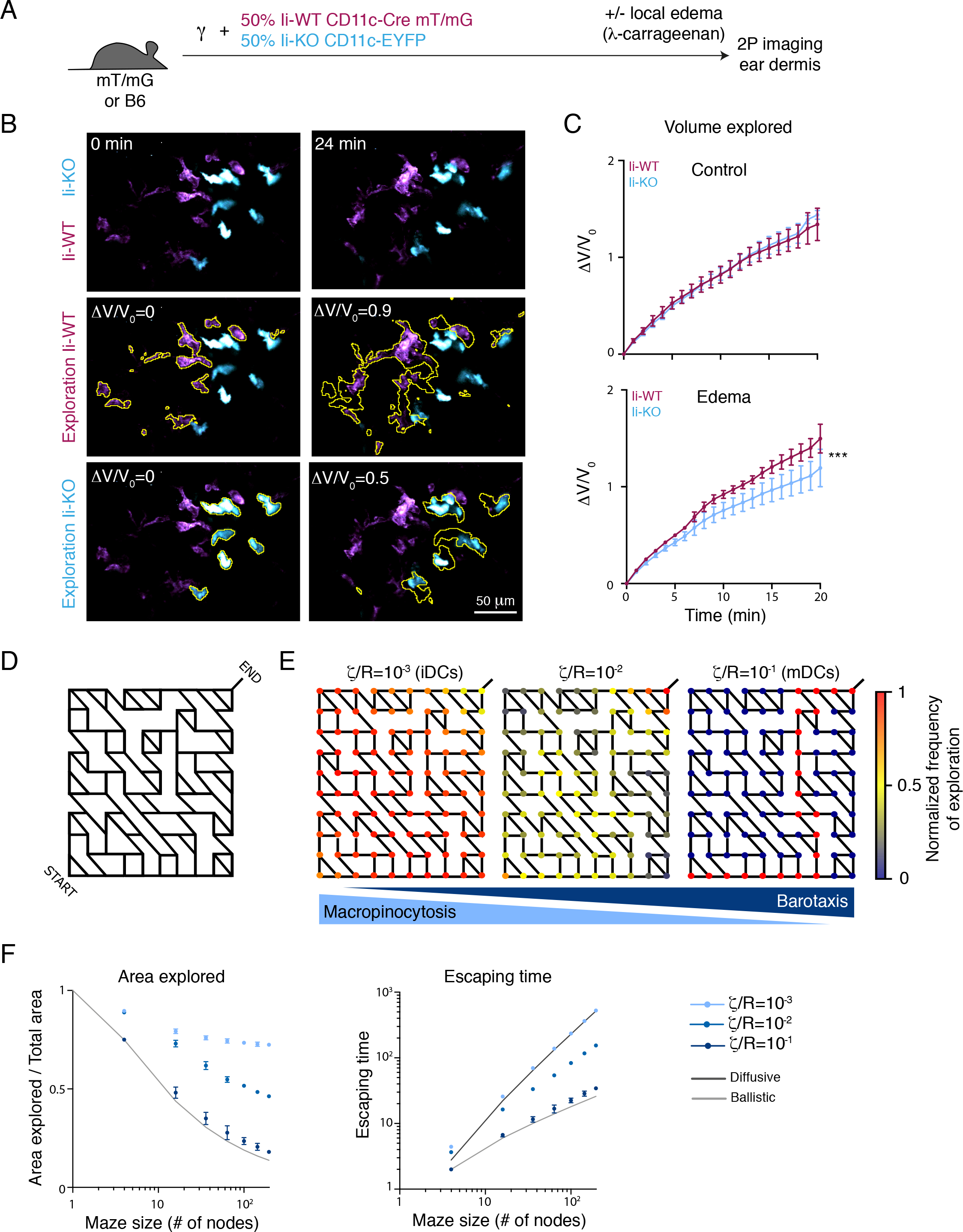
Macropinocytosis favors exploration, while barotaxis contributes to guidance. A. Experimental set-up for intravital imaging of dermal DCs in the ear, in the absence or in the presence of an edema. B. Time lapse images of ear dermis in local edema conditions. Ii-WT cells are in magenta, Ii-KO cells in cyan. The space explored is highlighted with a yellow dotted outline. The volume explored, normalized to the initial volume (ΔV/V_0_), is indicated on the images. C. The volume explored, normalized to the initial volume (ΔV/V_0_) is graphed over time. Data are averaged from 4 independent experiments. D. Design of the single exit maze used for simulations. E. Stochastic cell migration in random mazes has been simulated for different values of cell permeability to hydraulic resistance ratio. The panels represent the stochastic paths toward the exit followed by the simulated cells. The color code labels the normalized frequency of visiting a given node in the maze. F. Relative area explored and escaping time are graphed against the size of the maze. Solid curves correspond to the theoretical predictions for purely diffusive and ballistic migration modes.

Conversely, in mature DCs, which are no more exploratory but need to reach lymph nodes rapidly and efficiently, barotaxis might be beneficial. To address this question, we took advantage of our theoretical model by analyzing the stochastic dynamics of simulated cells in mazes whose nodes are interconnected randomly to only three nearest neighbors and whose edges have identical HR (**Fig. 5D**). The starting position in the maze is at the bottom left corner and the only exit is located at the opposite corner. Simulated cells visit multiple nodes before reaching the exit, and in each of them they encounter a Y-shaped bifurcation. Importantly, at each bifurcation, the path of lowest HR is precisely the shortest path to the exit. We observed two qualitatively different modes of exploration controlled by the ratio ζ/R between cell-fluid resistance and hydraulic resistance (See Supplementary Information). Highly permeable simulated cells (i.e. weakly barotactic, highly macropinocytic, ζ/R=10^−3^) chose randomly between both outgoing paths at every node. As a result, they tended to explore all accessible nodes in the random maze before exiting, performing a purely diffusive motion through the network (**Fig. 5E**, left). Impermeable simulated cells (i.e. strongly barotactic, weakly macropinocytic, ζ/R=10^−1^) were able to follow the path of lowest resistance, and therefore the shortest path to exit among all possible configurations (**Fig. 5E**, right). Explored area (mean area explored over the total network area) and escaping time (mean time to escape the maze) were quantified for both types of simulated cells (**Fig. 5E**). This showed that non barotactic cells were the most efficient at fully exploring the network, while barotactic cells, which selected the path with the least resistance at each node, were able to identify the shortest route across the random maze. Similar results were obtained for simulation in mazes presenting two exits (**Fig. S5**). This capacity to avoid dead-ends or longer paths might help mature dendritic cells to navigate the complex networks such as peripheral lymph vessels. Consistent with this finding, we had reported that CatS KO DCs, which do not down-regulate macropinocytosis as they mature, were defective when migrating from peripheral tissues to lymph nodes (Faure-Andre et al., 2008). Altogether, our experimental and model results suggest that modulation of barotaxis by macropinocytosis in dendritic cells might dose how much their physical environment affects their migration, to adapt their trajectories to their function as immature and mature cells.

In summary, we present here an approach that combines physical modeling and experiments to unravel the mechanisms underlying barotaxis of cells migrating in confinement. Our data show that barotaxis can be simply explained by a small asymmetry in the resisting forces due to HR, amplified by the acto-myosin cytoskeleton. We also found that, in iDCs, macropinocytosis cancels barotaxis, by increasing the permeability of cells to the extracellular fluid. This helps iDCs to explore larger territories than non-macropinocytic cells, which might facilitate their tissue sampling function. Upon microbial sensing, DCs mature and down-regulate macropinocytosis, thus regaining a barotactic behavior. Guidance by differences in HR might help mDCs avoiding dead ends while migrating to lymph nodes. This study therefore proposes a novel and unexpected role for macropinocytosis in modulating to which degree cell trajectories are biased by physical cues. It also suggests that HR might be an important cue for immune cell migration through tissues. How these physical cues interact with the chemical cues formed by chemokines gradients *in vivo* shall now be addressed.

## Acknowledgements

We acknowledge the Cell and Tissue Imaging Facility of the Institut Curie (PICT), a member of the France BioImaging national Infrastructure (ANR-10-INSB-04) and the Curie animal facility. We thank C. Hivroz and M. Bretou for insightful comments on the manuscript. We thank Z. Garcia, J. Postat and C. Grandjean from the Bousso lab for technical help. This work was supported by grants from Fondation pour la Recherche Médicale (FRM SPF20140129479) and Association pour la Recherche contre le Cancer (ARC-PDF20140601095) to HDM, the DCBIOL Labex (ANR-10-IDEX-0001-02-PSL and ANR-11-LABX-0043) to A.M.L.-D., as well as the ANR (PhyMax), the Fondation pour la Recherche Médicale and the Institut National du Cancer to A.-M.L.-D., M.P. and R.V.

## Author contributions

Conceptualization: HDM, CBM, RA, RV, MP, AMLD. Methodology: HDM, CBM, RA, PB, JFJ, RV, MP, AMLD. Investigation: HDM, RA, ZA, CBM. Formal analysis: HDM, CBM, ZA, MM. Resources: PB. Writing: HDM, CBM, RV, MP, AMLD. Visualization: HDM, CBM, MM. Supervision: RV, MP, AMLD.

## Materials and methods

### Cells

#### Dendritic cells

Bone-marrow derived dendritic cells (DCs) were obtained by culturing bone marrow cells for 10-11 days in IMDM medium supplemented with fetal calf serum (FCS, 10%), glutamine (20 mM), penicillin-streptomycin (100 U/mL), ß-mercaptoethanol (50 µM) and granulocyte-macrophage colony-stimulating factor (50 ng/mL)-containing supernatant obtained from transfected J558 cells, as previously described (Faure-Andre et al., 2008; Vargas et al., 2014). Immature DCs (iDCs) were collected by gentle recovery of semi-adherent cells. Mature DCs were obtained by 30 min treatment of iDCs with lipopolysaccharide (100 ng/mL) followed by careful washing with complete medium, and used 6-16h post-treatment, as previously described (Vargas et al., 2016).

#### HL60 cells

HL60 cells were cultured in RPMI-Glutamax completed with FCS (10%), penicillin-streptomycin (100 U/mL) and HEPES (25 mM). They were differentiated in neutrophils by adding DMSO (1.3%) to the culture medium for 5-6 days, as previously described (Millius and Weiner, 2010; Thiam et al., 2016).

### Mice

C57BL6/J mice were purchased from Charles River. LifeAct-GFP, Ii-KO, CatS-KO, CD11c-Cre Myosin-IIA-Flox, mT/mG (Muzumdar et al., 2007) and CD11c-Cre mT/mG mice were bred in our animal facilities. Bone marrow was collected from 8-week-old mice to generate DCs as described above. Bone marrow chimeric mice were generated by lethal γ-irradiation (9 Gy) of C57BL6/J or mT/mG recipient male mice, which were reconstituted 2 h later with a mixture of 50% CD11c-EYFP Ii-KO male bone marrow cells and 50% of CD11c-Cre^+^ mTmG^+^ Ii-WT male bone marrow cells. The chimeras were let 8-10 weeks to recover before being used for intravital imaging. All animal experiments were performed in accordance to the European and French Regulation for the Protection of Vertebrate Animals used for Experimental and other Scientific purposes (Directive 2010/63; French Decree 2013-118) and benefited from advices from the Animal Welfare Body of the Institut Curie Research Center and of the Institut Pasteur.

### Reagents

Fibronectin was purchased from Sigma and used at 10 µg/mL for coating microchannels. EIPA was purchased from Sigma and used at 50 µM (DMSO was used as a control). Rottlerin was purchased from Santa Cruz and used at 3 µM (DMSO was used as a control). CK666 was purchased from Tocris and used at 25 µM (DMSO was used as a control). Smifh2 was purchased from Tocris R&D Systems and used at 25 µM (DMSO was used as a control). Dextran-AlexaFluor 647 (10,000MW, Anionic, Fixable) was purchased from ThermoFisher and used at 4 mM. Lucifer Yellow (CH, Lithium Salt) was purchased from Fisher Scientific and used at 400 µM. *λ*-carrageenan was purchased from sigma and used resuspended at 1% in saline solution.

### Microbifurcation experiments and analysis

#### Microfabrication and imaging

Bifurcations were designed to exhibit different hydraulic resistances (HR) on each side, with a (R+ΔR)/R of 1, 5, 20 or infinite (dead-end). One additional design was made to compare high HR (same design as 20) to infinite HR (dead-end) (See **Fig. 1**) Two sizes of channels were generated: small channels were 6.5 µm wide and 2.8 µm high (thus had a cross-section of 18 µm^2^), large channels were 8.4 µm wide and 3.9 µm high (thus had a cross-section of 33 µm^2^). Microchannels were prepared as described previously (Faure-Andre et al., 2008) in polydimethylsiloxane (PDMS RTV 615, from Neyco), coated with Fibronectin for 1 h and washed with PBS before loading the cells in complete medium. Cells in channels were imaged for 6-12 h using an epifluorescence Nikon TiE video-microscope equipped with a cooled CCD camera (HQ2, Photometrics), using a 10X or a 20X objective.

#### Analysis

Image processing and analysis was performed using Fiji software (Schindelin et al., 2012). To quantify cell bias in bifurcations, only cells entering bifurcations devoid of any other cells were taken into account. Then, cells choosing the low resistance (or left) side of bifurcations (N_LR_) and the ones choosing the high resistance (or right) side of bifurcations (N_HR_) were counted manually. Bias was graphed as (N_LR_ − N_HR_)/(N_LR_ + N_HR_), so that a bias of 0 corresponds to 50% of cells choosing each side, and a bias of 1 corresponds to 100% of cells choosing low resistance (or left) side. Analysis of cell and actin dynamics in bifurcation was achieved using an homemade macro. Briefly, movies of individual cells passing bifurcation were cropped and rotated according to their channel direction. For each movie, the bifurcation position was manually selected on the transmission channel and 3 regions of interest were defined: before bifurcation, losing arm side and winning arm side. Then a mask of the migrating cell was obtained by thresholding on the GFP channel, and Analyze Particles function was used to get the cell position and instantaneous velocity at each time point. Additionally, the area of cell mask was measure for each time point in the 3 regions of interest. Thus we were able to define when the cell entered the bifurcation, when it chose the path to follow, and when it went out of the bifurcation. For averaging, time was normalized to take into account cell speed with 0 being the 1^st^ time point the cell entered the bifurcation and 1 being the last time point before it exited it. Then time was centered around 0 for retraction time.

### Macropinocytosis in microbifurcation assay and analysis

LifeAct-GFP DCs were imaged in microbifurcations filled or not with Dextran-Alexa647. Cell speed was quantified as described above. Cells were classified as positive for actin at the front if actin accumulation was clearly visible in front of the cell in at least one of the 3 time points before the bifurcation (for before choice) or in at least one of the 6 time points after the bifurcation (for after choice). Similarly, protrusion/retraction activity was quantified as at least one obvious event of protrusion and retraction in the 3 or 6 time points before or after bifurcation, resp.

### 2P imaging of the ear dermis and analysis

#### Intravital imaging

Two-photon imaging of the ear dermis was performed as previously described (Filipe-Santos et al., 2009). In brief, mice were anesthetized and placed on a custom-designed heated stage and a coverslip sealed to a surrounding parafilm blanket was placed on the ear, to immerge a heated 25X/1.05 NA dipping objective (Olympus). Imaging was performed using an upright FVMPE-RS microscope (Olympus). Multiphoton excitation was provided by an Insight DS + Dual laser (Spectra-Physics) tuned at 950 nm. Emitted fluorescence was split with 520, 562 and 506 nm dichroic mirrors and passed through 593/40 (mTom if present) and 542/27 (YFP) filters to nondescanned detectors (Olympus) and 483/32 (collagen by second harmonic generation) and 520/35 (GFP) filters to GASP detectors (Olympus). Typically, images from about 10 z planes, spaced 4 µm were collected every minute for up to one hour. Edema was induced by subcutaneous injection of 10 µL of *λ*-carrageenan (1% in saline). Imaging was performed 2 h after injection, around the site of injection.

#### Analysis

Image processing and analysis was performed using Fiji software (Schindelin et al., 2012). Two-photon Zstack movies were first registered to correct the drift. Then, for the analysis, some “clean/transparent” volumes were chosen and cropped from those movies. Each cropped sub-movies were converted to RGB and Color Threshold function was apply to obtain a 3D mask of GFP cells or YFP cells. Hue was set to 0-44 for YFP and 45-255 for GFP and the brightness parameter was chosen for different depth in tissue and interpolated for the other Z planes. After obtaining 3D masks movies, we generated exploration masks: for the time n, the 3D exploration mask was define as the merge between all 3D masks starting from time 0 to time n. Thus, by measuring the volume of the exploration mask upon the time (V) and normalizing it by the initial volume (V_0_, total volume of GFP or YFP cells at time 0), we obtained a measurement indicating the ability to explore tissue for both cell types. Volume explored are plotted as ΔV/V_0_ = (V-V_0_)/V_0_.

#### Statistical analysis

All statistical analysis has been made with GraphPad Prism software (v6).

## Figure legends

**Figure S1:**
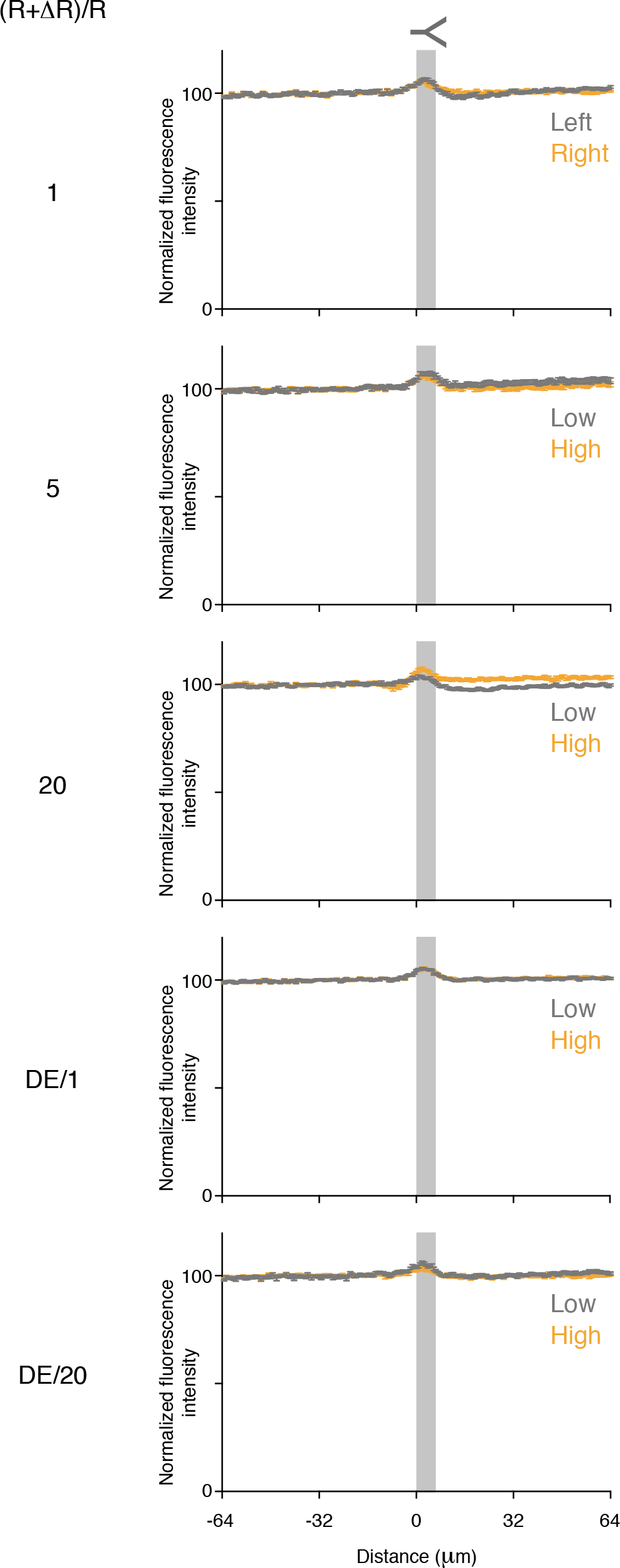
Microbifurcations are symmetrically filled regardless of hydraulic resistance asymmetry. Microbifurcations were filled with Lucifer-Yellow. Fluorescence profiles from a line along the center of the channel are plotted for both arms. The position of the bifurcation is indicated in grey. Note that there is no difference between both arms.

**Movie S1: Examples of iDCs, migrating in small channels presenting three types of bifurcation:** symmetric ((R+ΔR)/R=1, left), weakly asymmetric ((R+ΔR)/R=5, middle) or strongly asymmetric (DE/1, right).

**Figure S2:**
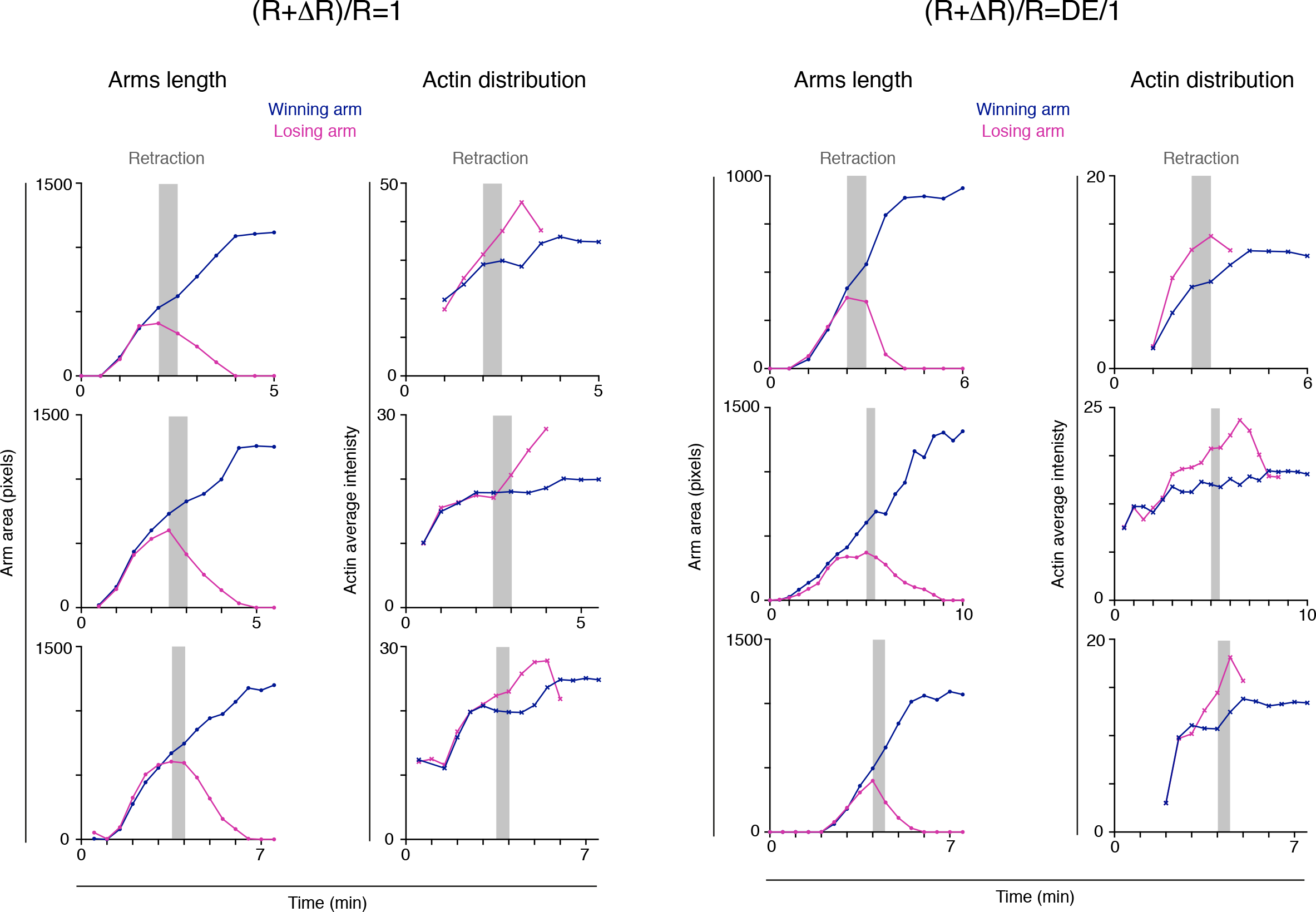
Example of arms lengths and actin distribution in bifurcations for single DCs.

**Movie S2: Actin distribution in simulated cell (left) and iDC (right) migrating in symmetric bifurcations.**

**Figure S3:**
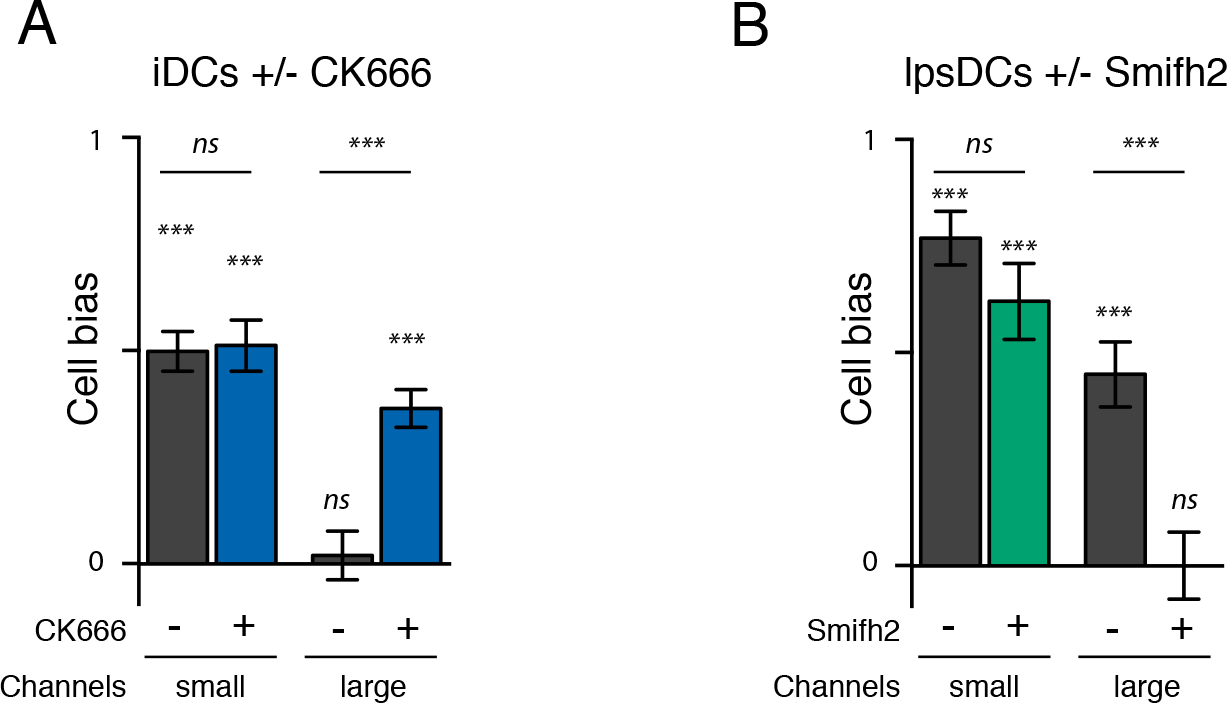
Modulation of macropinocytosis controls barotaxis. A. Bias of iDCs treated or not with the inhibitor of Arp2/3 CK666 to inhibit macropinocytosis (Vargas et al., 2016) are graphed. B. Bias of lpsDCs treated or not with the inhibitor of formins Smifh2 to restore macropinocytosis (Vargas et al., 2016) are plotted. Statistics above graph bars indicate difference to the value 0 (corresponding to no bias) calculated by a one sample t-test. Statistics between graph bars indicate difference between conditions calculated by a Mann-Withney test. ***: p-value < 0.001; **: p-value < 0.01; *: p-value < 0.05; *ns*: non significant. Data are pooled from at least 3 independent experiments.

**Figure S4:**
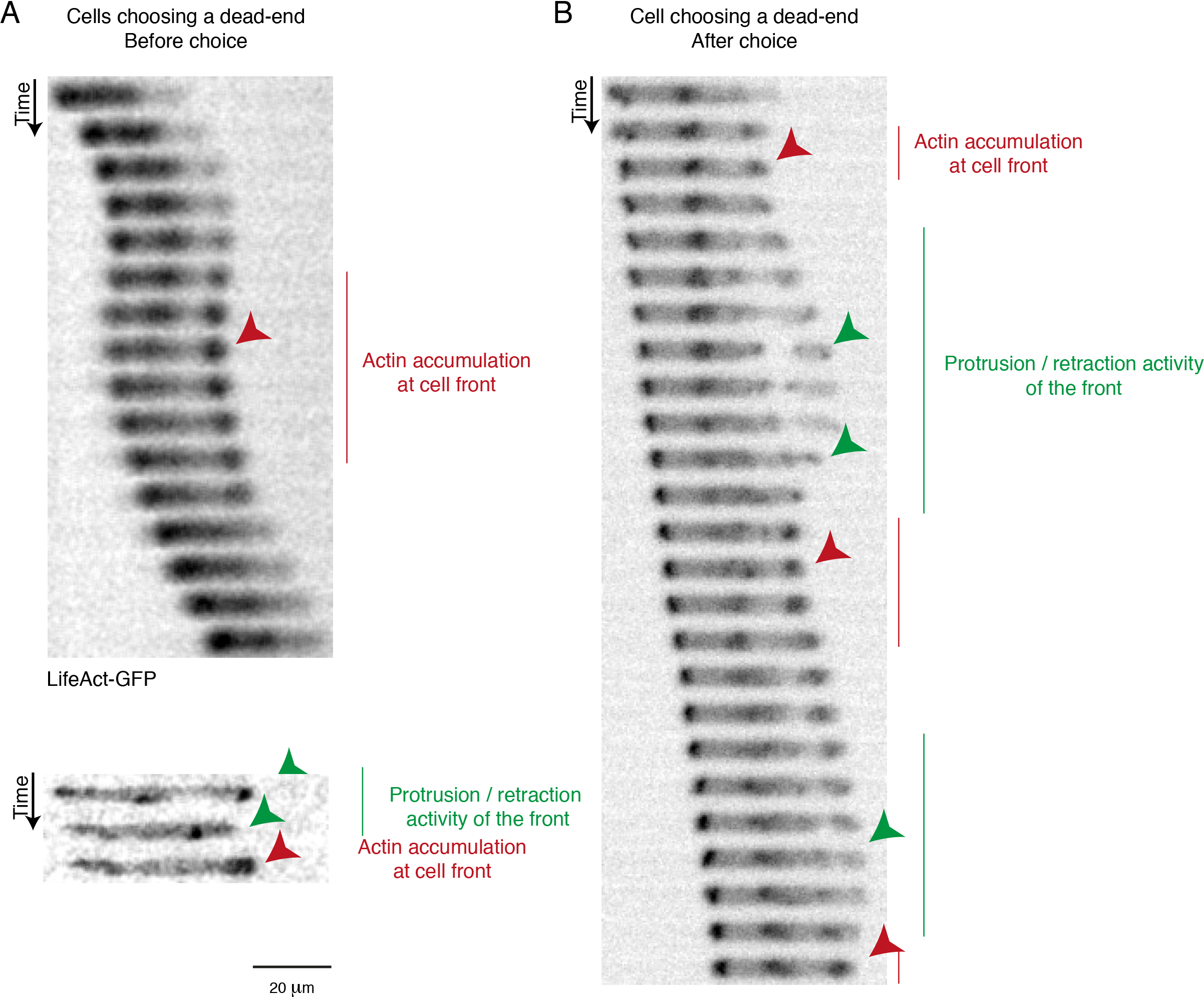
Examples of LifeAct-GFP distribution (inverted LUT) in single cells choosing dead-ends in small channels, showing actin accumulation at the front (red) and / or protrusion / retraction activity (green).

**Movie S3: Examples of iDCs migrating in open-ends or dead-ends, in small or large channels.** Green: LifeAct-GFP. Red: Dextran-AlexaFluor 647.

**Figure S5:**
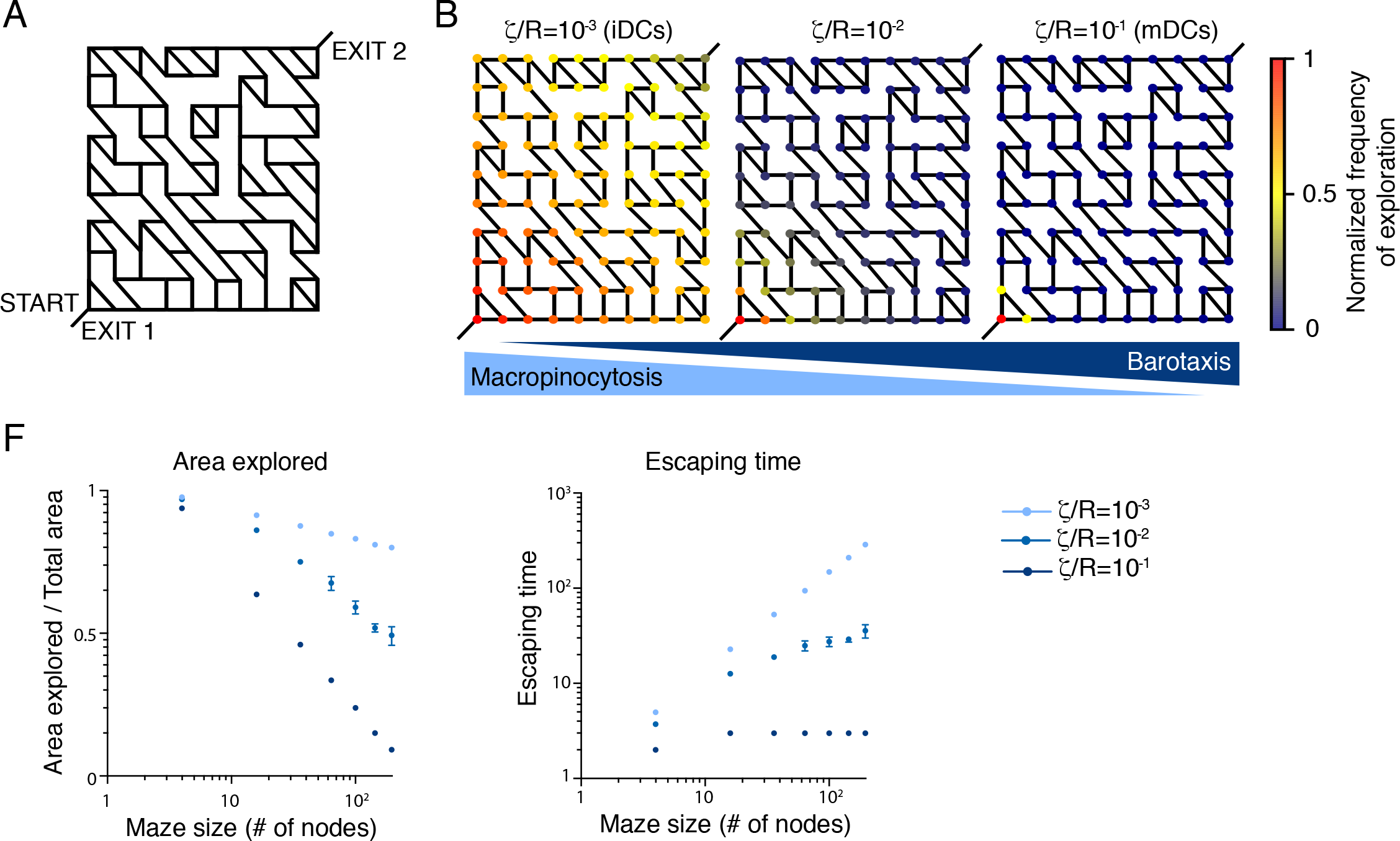
Exploration and escaping time of simulated cells in mazes with two exits. A. Design of the double exit maze used for simulations. B. Stochastic cell migration in random mazes has been simulated for different values of cell permeability to hydraulic resistance ratio. The panels represent the stochastic paths toward the exit followed by the simulated cells. The color code labels the normalized frequency of visiting a given node in the maze. C. Relative area explored and escaping time are graphed against the size of the maze.

**Movie S4: Space exploration by iDCs migrating at steady-state in the ear dermis.** Magenta: Ii-WT cells. Cyan: Ii-KO cells. Yellow outline: cumulated space explored.

**Movie S5: Space exploration by iDCs migrating after local edema induction in the ear dermis.** Magenta: Ii-WT cells. Cyan: Ii-KO cells. Yellow outline: cumulated space explored.

